# Application of Aligned-UMAP to longitudinal biomedical studies

**DOI:** 10.1101/2022.12.12.518225

**Authors:** Anant Dadu, Vipul K. Satone, Rachneet Kaur, Mathew J. Koretsky, Hirotaka Iwaki, Yue A. Qi, Daniel M. Ramos, Brian Avants, Jacob Hesterman, Roger Gunn, Mark R. Cookson, Michael E. Ward, Andrew B Singleton, Roy H Campbell, Mike A Nalls, Faraz Faghri

## Abstract

Longitudinal multi-dimensional biological datasets are ubiquitous and highly abundant. These datasets are essential to understanding disease progression, identifying subtypes, and drug discovery. Discovering meaningful patterns or disease pathophysiologies in these datasets is challenging due to their high dimensionality, making it difficult to visualize hidden patterns. Several methods have been developed for dimensionality reduction, but they are limited to cross-sectional datasets. Recently proposed Aligned-UMAP, an extension of the UMAP algorithm, can visualize high-dimensional longitudinal datasets. In this work, we applied Aligned-UMAP on a broad spectrum of clinical, imaging, proteomics, and single-cell datasets. Aligned-UMAP reveals time-dependent hidden patterns when color-coded with the metadata. We found that the algorithm parameters also play a crucial role and must be tuned carefully to utilize the algorithm’s potential fully.

Altogether, based on its ease of use and our evaluation of its performance on different modalities, we anticipate that Aligned-UMAP will be a valuable tool for the biomedical community. We also believe our benchmarking study becomes more important as more and more high-dimensional longitudinal data in biomedical research becomes available.

**Highlights:**

- explored the utility of Aligned-UMAP in longitudinal biomedical datasets
- offer insights on optimal uses for the technique
- provide recommendations for best practices

**In Brief:** High-dimensional longitudinal data is prevalent yet understudied in biological literature. High-dimensional data analysis starts with projecting the data to low dimensions to visualize and understand the underlying data structure. Though few methods are available for visualizing high dimensional longitudinal data, they are not studied extensively in real-world biological datasets. A recently developed nonlinear dimensionality reduction technique, Aligned-UMAP, analyzes sequential data. Here, we give an overview of applications of Aligned-UMAP on various biomedical datasets. We further provide recommendations for best practices and offer insights on optimal uses for the technique.

## Introduction

Visualizing large-scale, high-dimensional datasets is the starting step for any data exploratory analysis. Visualizing data is particularly useful for the biological community, where researchers rely on hypothesis-free data-driven analytics to gain essential insights and observe meaningful patterns from the data. The standard way of visualizing high-dimensional data is to project the data into low-dimensional space, typically 2D or 3D, while preserving local and global relationships. This transformation is called dimension reduction and belongs to the unsupervised machine learning algorithms class. The lower-dimensional data space can guide us in various tasks, such as identifying clusters, sub-structures, and outliers, detecting batch effects, and quality control measures to perform reliable and accurate downstream analyses.

In contrast to traditional methods for dimensionality reduction—for example, principal component analysis (PCA) (Jolliffe and Cadima 2016) — Uniform Manifold Approximation and Projection (UMAP) (McInnes, Healy, and Melville 2018) learns a nonlinear embedding of the original space by optimizing the embedding coordinates of individual data points using iterative algorithms. It aims to accurately preserve the original local neighborhood of each data point in the visualization. Because of the expressiveness of nonlinear embeddings, UMAP is well regarded for its state-of-the-art empirical performance at elucidating sophisticated manifold structures. The biomedical community widely adopts UMAP for multiple studies ranging from Single Cell RNAseq data (Becht et al. 2018)to Genetics (Diaz-Papkovich, Anderson-Trocmé, and Gravel 2021) or complex clinical symptoms (Becht et al. 2018)to depict exciting patterns from the data. In these use cases, UMAP is explored on datasets assuming that all samples in the dataset are independent.

Despite the prevalence of non-independent high-dimensional biological datasets, the application of UMAP in this area is little explored. This non-independence effect can occur from measurements at different time intervals, age, or other discrete/continuous variables. There are various longitudinal datasets of different modalities such as Clinical Symptoms, Magnetic Resonance Imaging (MRI), Electronic Health Records (EHR), Electroencephalography (EEG) for sleep monitoring, Electrocardiogram (ECG) data, etc. Since UMAP is a stochastic algorithm, different runs with the same hyperparameters can yield different results; therefore, extension to longitudinal datasets is not straightforward, unlike traditional algorithms such as PCA. Aligned-UMAP is a recently introduced dimensionality reduction approach for temporal data. It is based on the UMAP (McInnes, Healy, and Melville 2018)and MAPPER (Singh et al.2007)algorithms. MAPPER is a well-known topological data analysis method that successfully studies temporal, unbiased transcriptional regulation patterns (Rizvi et al. 2017). Aligned-UMAP imposes time constraints in the low dimensional embeddings, thereby controlling the stochasticity of its cross-sectional counterpart along the longitudinal axis. TimeCluster (Ali et al. 2019) is another approach that reduces the dimensionality of time-series data. Though it is possible to discover clusters with similar trajectories using TimeCluster, their intrinsic longitudinal variation cannot be observed. Further, it requires data availability for every time instance, making it less applicable for most biological datasets.

In this work, we deep dive into the applications of Aligned-UMAP on various longitudinal biological datasets. We applied the algorithm to clinical data, brain images, longitudinal proteomic data, EHR, and ECG datasets. We demonstrated its utility for researchers to identify exciting patterns and trajectories within enormous data sets. Secondly, we show the effect of different parameters of Aligned-UMAP on the lower dimension space. We also performed computation time analysis with varying datasets as a factor of the number of CPU cores. Furthermore, we deployed an interactive data visualization tool for reproducibility and transparency, motivated by open science. A deeper investigation of observed patterns could reveal more detailed, meaningful information, which is out of the scope of this work.

## Results

### Overview of the Aligned-UMAP method

#### Uniform Manifold Approximation and Projection

UMAP is a dimensionality reduction method that learns a non-linear low-dimensional embedding of the original high-dimensional space. UMAP has solid theoretical foundations based on manifold theory and preserves both the local and global structure better than other popular techniques such as t-SNE. UMAP is a graph-based dimensionality reduction method. It has two phases—first, computation of a weighted nearest-neighbor graph from the high-dimensional dataset. In the second phase, a low-dimensional layout is computed by optimizing the objective function that preserves desired characteristics of this nearest-neighbor graph. The algorithm is computationally efficient with the time order of sample size. It is the superior run time performance of UMAP that makes it very popular among the dimensionality reduction methods.

#### Aligned Uniform Manifold Approximation and Projection

Aligned-UMAP is a recently introduced dimensionality reduction approach for temporal data. The trivial way of performing dimensionality reduction on longitudinal data is to apply UMAP independently at different time steps and align the embedding using a Procrustes transformation on related points. However, Aligned-UMAP optimizes both embeddings simultaneously using a regularizer term to provide better alignments in general. The MAPPER algorithm is used to get the regularizer term which enforces the constraint on how far related points can take different locations in embeddings at multiple time points. Further details for the algorithm can be found on the UMAP documentation website ^1^. **Fig. 1** shows the pipeline of our analysis workflow.

**Fig 1.**
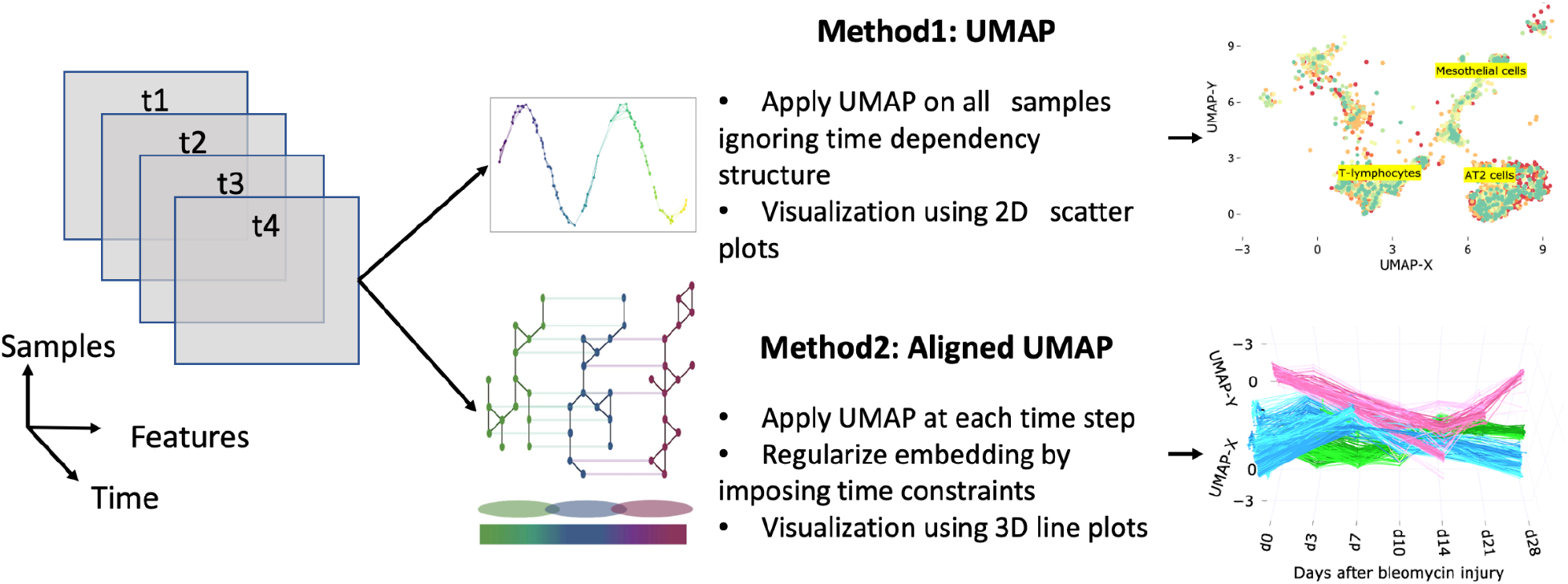
The workflow of analysis and model development.

### Software Output and Reproducibility

A demo of the Aligned-UMAP visualization is available at https://share.streamlit.io/anant-dadu/alignedumap-biomedicaldata. The data analysis pipeline for this work was performed in Python 3.8 using open-source libraries (numpy, pandas, plotly, umap). Our code is publicly available at https://github.com/NIH-CARD/AlignedUMAP-BiomedicalData to facilitate replication and future expansion of our work. The repository is well documented and includes a description of the data pre-processing, statistical, and machine learning analysis used in this study.

### Visualizing high-dimensional longitudinal data

We study Aligned-UMAP in a wide range of biomedical datasets from multiple data modalities. Table 1 shows the statistics of various datasets, with the count of samples ranging from approximately 500 to 21,000. These datasets vary in both the number of time sequences and the number of features available. For every visualization, each representative point becomes a thread through the time-axis as their relative position changes in the low-dimensional space.

**Table 1.** Datasets overview and statistics.

| Dataset | Modality | # samples | # features | # time sequences |
| --- | --- | --- | --- | --- |
| PPMI clinical data<sup>2</sup> | Clinical Assessment | 476 | 122 | 6 |
| ADNI clinical data<sup>3</sup> | Clinical Assessment | 435 | 78 | 4 |
| PPMI-ADNI T1 MRI | MRI T1 Imaging | 2,836 | 406 | 52 |
| MIMIC-III<sup>4</sup> (Johnson et al. 2016) | EHR | 36,675 | 64 | 6 |
| Longitudinal Proteomic COVID-19 | Proteomics | 383 | 1,463 | 3 |

<sup>1</sup> https://umap-learn.readthedocs.io/en/latest/aligned_umap_basic_usage.html
<sup>2</sup> http://www.ppmi-info.org/
<sup>3</sup> https://adni.loni.usc.edu/
<sup>4</sup> https://physionet.org/content/mimiciii-demo/1.4

|  |  |  |  |  |
| --- | --- | --- | --- | --- |
| Longitudinal whole lung scRNA | scRNA | 10,111 | 21,767 | 7 |
| iPSC derived neurons | Proteomics | 18 | 4,959 | 6 |

#### Clinical data

In neurodegenerative diseases such as Alzheimer’s and Parkinson’s, the individual can manifest disease in various ways, often times prior to clinical diagnosis. We evaluate the Aligned-UMAP algorithm on the clinical assessment data from Alzheimer’s Disease Neuroimaging Initiative (ADNI) and Parkinson’s Progression Markers Initiative (PPMI) study cohorts. The ADNI study includes Alzheimer’s patients, mild cognitive impairment subjects, and elderly controls. PPMI study has subjects recently diagnosed with Parkinson’s Disease (PD) and healthy controls. These studies collect data for many clinical assessments related to movement and cognitive disability to monitor disease progression. All such measurements are recorded longitudinally at separate visits. The time duration of such visits can range from years to decades.

We preprocess the ADNI and PPMI cohort datasets following the strategy proposed in previous disease subtyping studies (Faghri et al.2018; Satone et al. 2020). UMAP and Aligned-UMAP successfully pulled together clusters corresponding to populations with similar disease progression (**Fig 2a** and **Fig 2b**). However, longitudinal differences got lost in the UMAP version due to its stochastic nature. Aligned-UMAP separates rapidly progressive PD from the healthy control group and demonstrates divergence of the rapid PD subgroup from healthy controls with aging (**Fig 2a**). Furthermore, Aligned-UMAP reveals distinct longitudinal courses for dementia and the healthy control group **(Fig. 2b**). We follow a continuum spectrum from lower progressive to high progressive subgroups for PD and Dementia subjects (**Supplementary Fig. 1**). These results suggest that Aligned-UMAP could be used as a hypothesis-generating tool to identify distinct subtypes based on disease progression. For instance, a particular subgroup shows rapid decline in clinical symptoms such as MDS-Unified Parkinson’s Disease Rating Scale (Goetz et al. 2008) or MoCA cognitive assessment (Nasreddine et al. 2005) as compared to healthy control and other subgroups.

**Fig 2.**
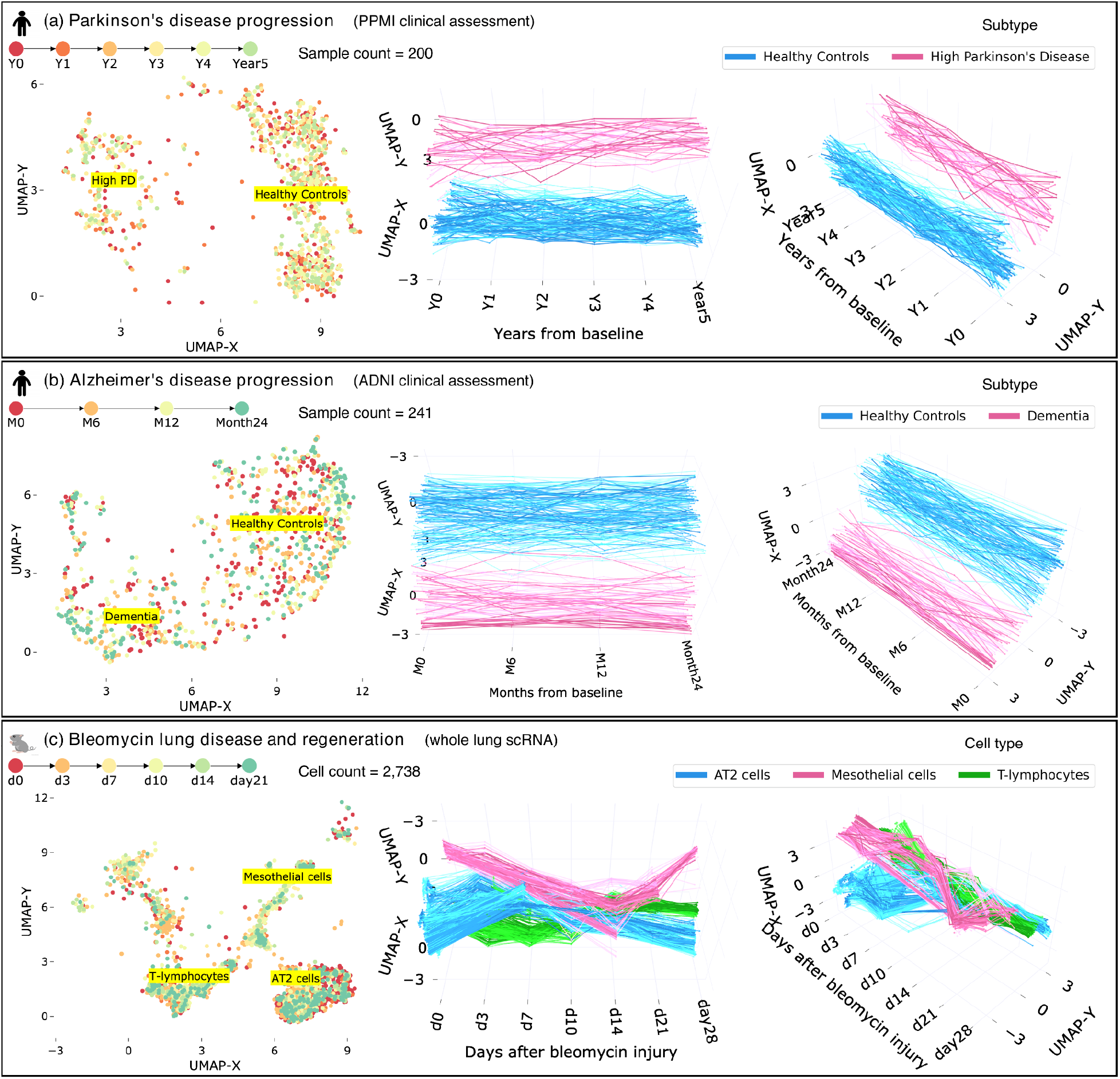
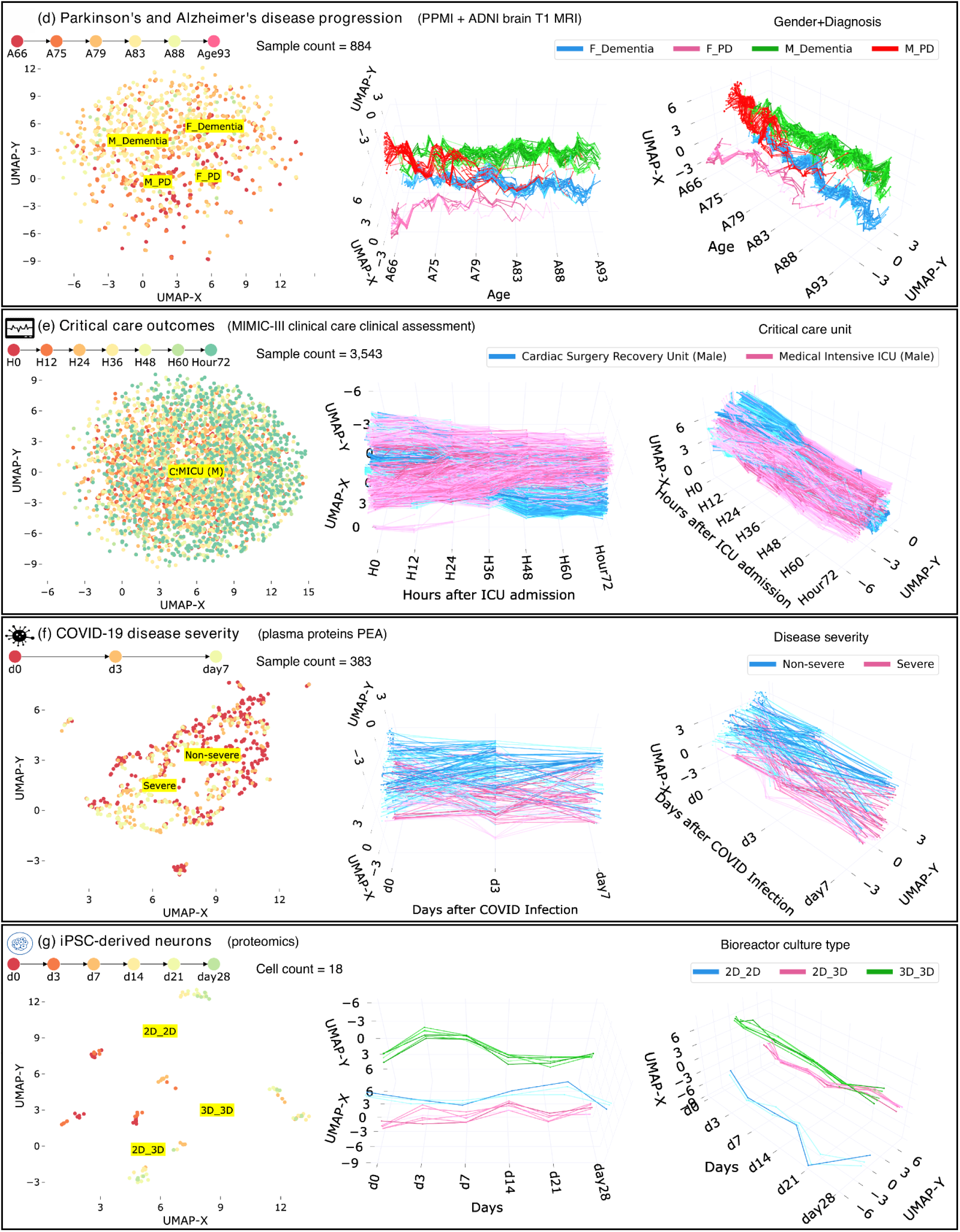
Low dimensional embeddings by UMAP and Aligned-UMAP dimensionality reduction algorithms on longitudinal biomedical datasets from multiple modalities. (a) Reveals the distinction between Parkinson’s Disease subjects (with rapid progressors) and Healthy Controls from 122 clinical measurements collected over five years from Parkinson’s Progression Markers Initiative (PPMI) study. Measures include Montreal Cognitive Assessment scores and MDS-Unified Parkinson’s Disease Rating Scale scores. (b) Show trajectories of Dementia and Healthy Control subjects on 78 clinical measurements collected over two years from the Alzheimer’s Disease Neuroimaging Initiative (ADNI) study. Measurements include Mini-Mental State Exam (MMSE) scores and Alzheimer’s Disease Assessment Scale–Cognitive Subscale (ADAS-COG) tests. (c) Aligned-UMAP trajectories show shifts in specific cell types (such as Mesothelial and AT2 cells) in gene expression space during the regeneration time course of mice having bleomycin lung injury. (d) Aligned-UMAP embeddings depict aging patterns for Dementia, and Parkinson’s disease patients, stratified by gender. (e) Unveils trajectories of the subject’s admitted in different critical care units of the MIMIC-III database. Measurements include vital signs such as blood pressure, oxygen levels, and ICD-9 diagnosis codes. (f) Embedding space depicts the severity of COVID-19 disease from 1,463 unique plasma proteins measured by proximity extension assay using the Olink platform. The cutoff at day 3 is visible because of data unavailability at day seven due to either patient recovery or deceased (g) Aligned-UMAP low dimensional space identified the cell culture environment of iPSC-derived neurons using longitudinal proteomic data for more than 8000 proteins. Note: We apply the Aligned-UMAP algorithm on the dataset having characteristics shown in Table 1. In this figure, we have demonstrated a subset of classes for better visualization purposes. For more detailed analysis, users can explore our public web application.

#### Whole lung scRNA Data

Single-cell transcriptomics (scRNA) using next-generation transcript sequencing (RNA-seq) has recently received much attention due to its ability to uncover cellular heterogeneity, cellular differentiation, and development mechanisms. UMAP has demonstrated its efficacy in analyzing single-cell datasets by identifying clusters of related cells. Modeling gene expression trajectories of different cell types have been successfully used to understand cell-cell communication routes in various chronic diseases such as lung disease and tumor cells (Strunz et al. 2020;Sharma et al. 2018). We evaluated Aligned-UMAP on whole lung scRNA data of mice undergoing regeneration after bleomycin-induced lung injury (Strunz et al. 2020). Transcriptomic profile of 29,297 cells was collected from six time points (day 3, 7, 10, 14, 21, and 28). We observe clusters of cell types showing different cellular dynamics through the regeneration process (**Fig. 2c**); Mesothelial cells show a spike at day 14 and start returning to their healthy state (day 0). Thereby, suggesting the role of mesothelial cells in bleomycin related lung injury. This way, we could extract hidden longitudinal patterns from high-dimensional time-series datasets using Aligned-UMAP.

#### Imaging Data

Imaging is a pervasive way of monitoring the disease progression of multiple disorders. We use the Advanced Normalization Tools (ANTs) pipeline(Tustison et al. 2021) to extract structural features such as the volume and area of different brain regions from the Magnetic Resonance Imaging (MRI) T1 image. Since the number of longitudinal images for each subject is scarce, we use the imaging features to model aging trajectories. To be precise, we relate images if they are observed at similar age groups instead of relating subjects based on their visits. Also, these relations are constrained by different diagnosis groups (i.e., Control, PD, or Dementia). **Fig. 2d** shows various aging courses based on the subject’s latest diagnosis and gender. We noticed a more rapid decline among female dementia cases versus male dementia cases around 80 years of age. It suggests the non-linear and distinct patterns of disease progression across groups within a disease. We observed distinct longitudinal trajectory patterns, which might be a possible way to monitor disease progression^5^.

#### EHR Data

Electronic Health Records (EHR) is a systematic collection of patients’ healthcare records in a digital format. EHR is adopted in many hospitals in the USA and UK (Adler-Milstein et al. 2017). We applied the Aligned-UMAP on the MIMIC-III Critical Care Database (Johnson et al. 2016), which consists of records of more than 40,000 patients in intensive care units (ICU) of the Beth Israel Deaconess Medical Center between 2001 and 2012. We preprocessed the dataset following the methodology proposed by (Lin et al. 2019). **Fig. 2 (e)** shows the lower-dimensional space on the MIMIC-III dataset on measurement recorded during the initial 72 hrs of entry to the Intensive Care Unit (ICU). We color the trajectories based on the type of critical care unit a patient stays in just before discharge from the hospital. We observe that UMAP could not recover time-related patterns; however, Aligned-UMAP segregates trajectories based on the patient’s critical care unit. This pattern reflects that it might be helpful to analyze ICU datasets stratified by their care unit and suggests that the quality of care in ICU units is highly variable.

#### COVID-19 Proteomics Data

Uncovering protein signatures associated with COVID-19 infection and severity can provide insights into its pathophysiology and immune dysfunction (Filbin et al. 2021). We utilized longitudinal proteomic data on 306 COVID-19 patients (Filbin et al. 2021). Aligned-UMAP has identified distinct trajectories for severe and non-severe patients over seven days **(Fig. 2f)**. We observed the participants exhibiting continued negative symptom trajectories at seven days belonging to more severe or longer COVID-19 infection.

#### iPSC-derived neurons Proteomics Data

Aligned-UMAP can be incorporated as a quality control measure for longitudinal data. We applied this approach to longitudinal proteomic profiling of the differentiation of iPSC (Induced Pluripotent Stem Cells) derived neurons cultured in different bioreactors (Reilly et al.2021). We could visualize distinct patterns of change for each cell line grouped by their culture environment, thereby identifying batch effects (**Fig. 2g**). We observed that the cell lines cultured only in the 2D bioreactor are hypervariable for almost all time points (till day 28). The cell line 2D_3D (day0-day3 2D culture, day4-day28 3D culture) tends to converge around day 14, and the cell line cultured in the 3D bioreactor tends to be more homogenous after around day 7.A tighter spread denotes a homogenous group.

## Discussion

### Observed Meaningful Patterns

Our work demonstrates that Aligned-UMAP could help us discover meaningful longitudinal patterns by color-coding them based on multiple known covariates. Our analysis finds that both UMAP and Aligned-UMAP help generate intuitive embeddings because of their ability to preserve the global structure. Additionally, Aligned-UMAP provides a view that highlights longitudinal structure by imposing time constraints in the embeddings, thereby controlling the stochasticity of its cross-sectional counterpart. We observe distinct trajectory patterns of the data from different modalities. Dementia and PD subtypes are delineated using clinical assessment measurements from the PPMI and ADNI study (**Fig. 2a, Fig. 2b**). Aligned-UMAP has also shown visually meaningful patterns on high-dimensional omics data such as proteomics (**Fig. 2f, Fig. 2g**) or single-cell transcriptomics data (**Fig. 2c**). Therefore, it is evident that Aligned-UMAP provides meaningful representations and is likely to be a valuable tool for researchers working on multivariate longitudinal datasets by preserving the global and local trends along the time axis.

### Points to remember

Based on our observations from this study, this approach promises to be useful in many other biomedical datasets. These datasets can vary in terms of data missingness, time sequences, or domain-specific variations that make it challenging to tune experimental settings. So, here we discuss key points that users should keep in mind while using Aligned-UMAP.

- **Data Missingness Effect:** The problem of missing data is prevalent in healthcare datasets and can interfere with the conclusions drawn from the data. Aligned-UMAP can handle data missingness across the longitudinal dimension by performing interpolation in low-dimensional space. Tensor decomposition based dimension reduction approaches cannot handle any data missingness (Ali et al. 2019). However, none of the dimension reduction approaches are designed to handle missingness for features measured cross-sectionally.
- **Aligned-UMAP Parameter Effect:** The number of neighbors and the minimum distance are two critical parameters affecting the lower-dimensional space using the UMAP algorithm. In Aligned-UMAP, the number of parameters can increase significantly. We can vary the UMAP parameters for each step to observe different trajectories. The two other alignment parameters, namely, *alignment window size* and *alignment regularizer*, are critical in visualizing the longitudinal trend that controls the volatility along the time axis. **Fig. 3** shows the effect of *alignment window size, alignment regularizer*, and the *number of neighbors* on the PPMI longitudinal dataset. Our web app also demonstrates the impact of these parameters on the lower-dimensional space.
- **Execution Time:** We analyze the execution time taken by both algorithms on multiple datasets and use their subsamples of different sizes. Further, to understand the algorithm’s scalability and parallelization, we executed it utilizing different numbers of cores (**Fig. 4**). Multiple core setup does not seem to improve run times of Aligned-UMAP in low data regimes, which may be attributed to inter-core synchronization overheads. However, significant improvements are observed on complete lung scRNA data with 16 cores (**Fig. 4a**). Compared to UMAP, Aligned-UMAP would require a larger dataset to have better parallelization on a multi-core machine (**Fig. 4b**).
- Stochastic models and reproducibility: Although Aligned-UMAP can handle stochasticity along the longitudinal axis, it still produces variable embeddings on different runs. Like UMAP, it uses randomness both to speed up approximation steps and to aid in solving optimization problems, thereby affecting the reproducibility of the lower dimensional space. However, UMAP and Aligned-UMAP provide relatively stable results when applied to large amounts of data. In the future, sophisticated approaches are required to ensure reproducibility.

**Fig 3.**
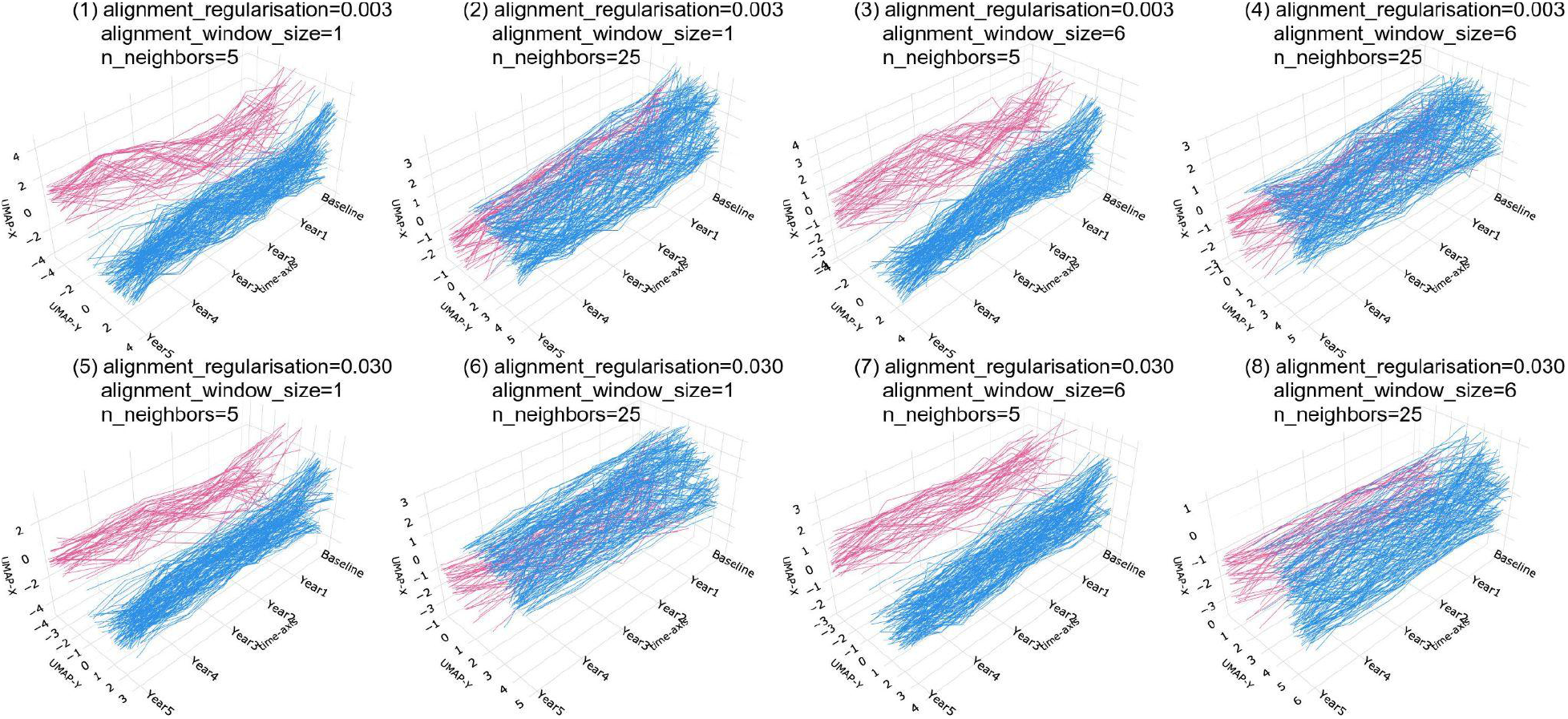
Effect of hyperparameters of Aligned-UMAP on the PPMI clinical dataset. The alignment regularization is varied for [0.003, 0.03], alignment window size from [1, 6] and number of neighbors from [5, 25]. We could observe that an increase in the number of neighbors increases the size of visible clusters (1, 2). Alignment regularization controls and alignment window size the volatility of trajectories. Higher values for alignment regularization will keep the related embeddings closer (1, 5), and alignment window size captures how far forward and backward across the datasets we look at when doing alignment (1, 3).

**Fig 4.**
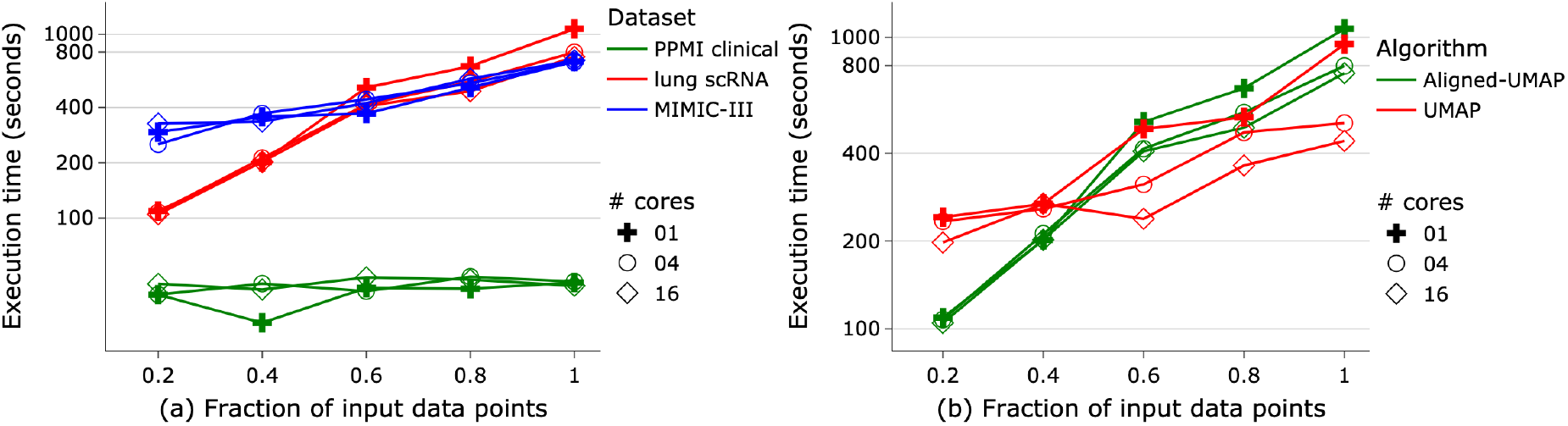
Execution time for input datasets of varying sizes (a) for Aligned-UMAP on multiple datasets (b) for Aligned-UMAP and UMAP on whole lung scRNA dataset. All the experiments are conducted on a 128 GB RAM machine utilizing a different number of cores (marker symbol).

## Future Work

The Aligned-UMAP algorithm is still in the developing phase. We discuss the plausible extensions of the algorithm that might be useful in a multitude of biomedical research datasets.

- **Clustering:** The dimensionality reduction method is a standard preprocessing step to utilize density-based clustering methods on the high-dimensional dataset. Dynamic time warping is the most common metric to cluster time-varying patterns using K-mean clustering. It will be interesting to evaluate multiple clustering approaches on longitudinal trajectories.
- **Semi-Supervised / Supervised:** Sometimes, we would like to incorporate target label information to project high-dimensional data to lower-dimensional space in dimensionality reduction. There are various reasons for supervised dimension reduction; First, to retain the internal structure of classes and have dense clusters. Secondly, to maintain the global structure. i.e., preservation of inter-relationships among the known classes. Finally, we can observe outliers or subjects that do not belong to either class using the semi-supervised learning approach. The extension of Aligned-UMAP for supervised/semi-supervised dimension reduction will be a part of future work.
- **Rare Events Detection:** UMAP algorithm supports the detection of outliers using the Local Outlier Factor(Breunig et al. 2000) algorithm. Identifying outliers from longitudinal trajectories generated by Aligned-UMAP will need further investigation.
- **Multimodal aspect:** In the biomedical domain, monitoring disease needs data from multiple modalities such as imaging, blood biomarkers, genetics, or multi-omics (Makarious et al. 2022). Current dimensionality reduction approaches are destined for the dataset from a single modality. The trivial way of incorporating multimodal data is to use vectorization, but it might not be the optimal solution to discover hidden patterns in the data. Therefore, evaluating and building new dimensionality reduction approaches for multimodal data analysis setup is required.
- **Interpretability**: It’s important to note that because UMAP and t-SNE both necessarily warp the high-dimensional shape of the data when projecting to lower dimensions, any given axis or distance in lower dimensions still isn’t directly interpretable in the way of techniques such as PCA.
- **Data frequency:** Since Aligned-UMAP creates a lower-dimensional space for every location, analyzing data collected at an extremely fine scale, such as ICU or ECG spectrograms, becomes expensive.

## Acknowledgments

We thank the patients and their families who contributed to this research. This research was supported in part by the Intramural Research Program of the National Institute on Aging (NIA) and National Institute of Neurological Disorders and Stroke (NINDS), both part of the National Institutes of Health, within the Department of Health and Human Services; project number ZIA AG000534, ZO1 AG000949, ZIA-NS003154, and the Michael J Fox Foundation. Data used in the preparation of this article were obtained from the Parkinson’s Progression Markers Initiative (PPMI) database (www.ppmi-info.org/data). For up-to-date information on the study, visit www.ppmi-info.org. PPMI – a public-private partnership – is funded by the Michael J. Fox Foundation for Parkinson’s Research and funding partners, including Abbvie, Avid Radiopharmaceuticals, Biogen Idec, Bristol-Myers Squibb, Covance, Eli Lilly & Co., F. Hoffman-La Roche, Ltd., GE Healthcare, Genentech, GlaxoSmithKline, Lundbeck, Merck, MesoScale, Piramal, Pfizer, and UCB. Data and biospecimens used in the preparation of this manuscript were obtained from the Parkinson’s Disease Biomarkers Program (PDBP) Consortium, part of the National Institute of Neurological Disorders and Stroke at the National Institutes of Health. Investigators include: Roger Albin, Roy Alcalay, Alberto Ascherio, DuBois Bowman, Alice Chen-Plotkin, Ted Dawson, Richard Dewey, Dwight German, Xuemei Huang, Rachel Saunders-Pullman, Liana Rosenthal, Clemens Scherzer, David Vaillancourt, Vladislav Petyuk, Andy West and Jing Zhang. The PDBP Investigators have not participated in reviewing the data analysis or content of the manuscript.

Data collection and sharing for this project was funded by the Alzheimer’s Disease Neuroimaging Initiative (ADNI) (National Institutes of Health Grant U01 AG024904) and DOD ADNI (Department of Defense award number W81XWH-12-2-0012). ADNI is funded by the National Institute on Aging, the National Institute of Biomedical Imaging and Bioengineering, and through generous contributions from the following: AbbVie, Alzheimer’s Association; Alzheimer’s Drug Discovery Foundation; Araclon Biotech; BioClinica, Inc.; Biogen; Bristol-Myers Squibb Company; CereSpir, Inc.; Cogstate; Eisai Inc.; Elan Pharmaceuticals, Inc.; Eli Lilly and Company; EuroImmun; F. Hoffmann-La Roche Ltd and its affiliated company Genentech, Inc.; Fujirebio; GE Healthcare; IXICO Ltd.; Janssen Alzheimer Immunotherapy Research & Development, LLC.; Johnson & Johnson Pharmaceutical Research & Development LLC.; Lumosity; Lundbeck; Merck & Co., Inc.; Meso Scale Diagnostics, LLC.; NeuroRx Research; Neurotrack Technologies; Novartis Pharmaceuticals Corporation; Pfizer Inc.; Piramal Imaging; Servier; Takeda Pharmaceutical Company; and Transition Therapeutics. The Canadian Institutes of Health Research is providing funds to support ADNI clinical sites in Canada. Private sector contributions are facilitated by the Foundation for the National Institutes of Health (www.fnih.org). The grantee organization is the Northern California Institute for Research and Education, and the study is coordinated by the Alzheimer’s Therapeutic Research Institute at the University of Southern California.

## Data Availability

The data used in this study was access-controlled from the Parkinson’s Progression Marker Initiative (PPMI, http://www.ppmi-info.org/) and the the Alzheimer’s Disease Neuroimaging Initiative (ADNI, https://adni.loni.usc.edu). and require individual sign-up to access the data. Electronic health records from MIMIC-III Critical Care Database were downloaded from https://physionet.org/content/mimiciii-demo/1.4. Bulk and scRNA-seq data from mice whole lung are available via the Gene Expression Omnibus with the accession code GSE141259. COVID-19 longitudinal proteomic data have been downloaded from http://dx.doi.org/10.17632/nf853r8xsj. Additionally, we have developed an interactive website [https://share.streamlit.io/anant-dadu/alignedumap-biomedicaldata] where researchers can investigate components of the predictive model and can investigate feature effects on a sample and cohort level.

## Author contributions

A.D., A.B.S., M.A.N., R.H.C., and F.F. contributed to the concept and design of the study. A.D., V.K.S., R.K. H.I., Y.A.Q., D.M.R., B.A., J.H., R.G., M.R.C., M.E.W., A.B.S., R.H.C, M.A.N., and F.F. were involved in the acquisition of data, data generation, and data cleaning. A.D., R.H.C., M.A.N., and F.F. did the analysis and interpretation of data. A.D., H.I., Y.A.Q., D.M.R., B.A., J.H., R.G., M.R.C., M.E.W., A.B.S., R.H.C, M.A.N., and F.F contributed to the drafting of the article and revising it critically.

## Code Availability

To facilitate replication and expansion of our work, we have made the notebook publicly available on GitHub at [https://github.com/NIH-CARD/AlignedUMAP-BiomedicalData]. It includes all code, figures, models, and supplements for this study. The code is part of the supplemental information; it includes the rendered Jupyter notebook with full step-by-step data preprocessing, statistical, and machine learning analysis.

## Footnotes

1 https://umap-learn.readthedocs.io/en/latest/aligned_umap_basic_usage.html

2 http://www.ppmi-info.org/

3 https://adni.loni.usc.edu/

4 https://physionet.org/content/mimiciii-demo/1.4

5 Further investigation of trajectory patterns is out of scope of this work.

## References

Adler-Milstein, Julia, A. Jay Holmgren, Peter Kralovec, Chantal Worzala, Talisha Searcy, and Vaishali Patel. 2017. “Electronic Health Record Adoption in US Hospitals: The Emergence of a Digital ‘Advanced Use’ Divide.” Journal of the American Medical Informatics Association: JAMIA 24 (6): 1142–48.

Ali, Mohammed, Mark W. Jones, Xianghua Xie, and Mark Williams. 2019. “TimeCluster: Dimension Reduction Applied to Temporal Data for Visual Analytics.” The Visual Computer 35 (6): 1013–26.

Becht, Etienne, Leland McInnes, John Healy, Charles-Antoine Dutertre, Immanuel W. H. Kwok, Lai Guan Ng, Florent Ginhoux, and Evan W. Newell. 2018. “Dimensionality Reduction for Visualizing Single-Cell Data Using UMAP.” Nature Biotechnology, December. https://doi.org/10.1038/nbt.4314.

Breunig, Markus M., Hans-Peter Kriegel, Raymond T. Ng, and Jörg Sander. 2000. “LOF: Identifying Density-Based Local Outliers.” SIGMOD Rec. 29 (2): 93–104.

Dadu, Anant, Vipul K. Satone, Rachneet Kaur, Sayed Hadi Hashemi, Hampton Leonard, Hirotaka Iwaki, Mary B. Makarious et al. “Identification and prediction of Parkinson’s disease subtypes and progression using machine learning in two cohorts.” bioRxiv (2022). https://doi.org/10.1101/2022.08.04.502846

Diaz-Papkovich, Alex, Luke Anderson-Trocmé, and Simon Gravel. 2021. “A Review of UMAP in Population Genetics.” Journal of Human Genetics 66 (1): 85–91.

Faghri, Faraz, Sayed Hadi Hashemi, Hampton Leonard, Sonja W. Scholz, Roy H. Campbell, Mike A. Nalls, and Andrew B. Singleton. 2018. “Predicting Onset, Progression, and Clinical Subtypes of Parkinson Disease Using Machine Learning.” bioRxiv. https://doi.org/10.1101/338913.

Faghri, Faraz, Fabian Brunn, Anant Dadu, Adriano Chiò, Andrea Calvo, Cristina Moglia, Antonio Canosa et al. “Identifying and predicting amyotrophic lateral sclerosis clinical subgroups: a population-based machine-learning study.” The Lancet Digital Health 4, no. 5 (2022): e359–e369.

Filbin, Michael R., Arnav Mehta, Alexis M. Schneider, Kyle R. Kays, Jamey R. Guess, Matteo Gentili, Bánk G. Fenyves, et al. 2021. “Longitudinal Proteomic Analysis of Severe COVID-19 Reveals Survival-Associated Signatures, Tissue-Specific Cell Death, and Cell-Cell Interactions.” Cell Reports. Medicine 2 (5): 100287.

Goetz, Christopher G., Barbara C. Tilley, Stephanie R. Shaftman, Glenn T. Stebbins, Stanley Fahn, Pablo Martinez-Martin, Werner Poewe, et al. 2008. “Movement Disorder Society-Sponsored Revision of the Unified Parkinson’s Disease Rating Scale (MDS-UPDRS): Scale Presentation and Clinimetric Testing Results.” Movement Disorders: Official Journal of the Movement Disorder Society 23 (15): 2129–70.

Johnson, Alistair E. W., Tom J. Pollard, Lu Shen, Li-Wei H. Lehman, Mengling Feng, Mohammad Ghassemi, Benjamin Moody, Peter Szolovits, Leo Anthony Celi, and Roger G. Mark. 2016. “MIMIC-III, a Freely Accessible Critical Care Database.” Scientific Data 3 (May): 160035.

Jolliffe, Ian T., and Jorge Cadima. 2016. “Principal Component Analysis: A Review and Recent Developments.” Philosophical Transactions. Series A, Mathematical, Physical, and Engineering Sciences 374 (2065):20150202.

Lin, Yu-Wei, Yuqian Zhou, Faraz Faghri, Michael J. Shaw, and Roy H. Campbell. 2019. “Analysis and Prediction of Unplanned Intensive Care Unit Readmission Using Recurrent Neural Networks with Long Short-Term Memory.” PloS One 14 (7): e0218942.

Makarious, Mary B., Hampton L. Leonard, Dan Vitale, Hirotaka Iwaki, Lana Sargent, Anant Dadu, Ivo Violich et al. “Multi-modality machine learning predicting Parkinson’s disease.” npj Parkinson’s Disease 8, no. 1 (2022): 1–13.

McInnes, Leland, John Healy, and James Melville. 2018. “UMAP: Uniform Manifold Approximation and Projection for Dimension Reduction.” arXiv [stat.ML]. arXiv. http://arxiv.org/abs/1802.03426.

Nasreddine, Ziad S., Natalie A. Phillips, Valérie Bédirian, Simon Charbonneau, Victor Whitehead, Isabelle Collin, Jeffrey L. Cummings, and Howard Chertkow. 2005. “The Montreal Cognitive Assessment, MoCA: A Brief Screening Tool for Mild Cognitive Impairment.” Journal of the American Geriatrics Society 53 (4): 695–99.

Reilly, Luke, Lirong Peng, Erika Lara, Daniel Ramos, Michael Fernandopulle, Caroline B. Pantazis, Julia Stadler, et al. 2021. “A Fully Automated FAIMS-DIA Proteomic Pipeline for High-Throughput Characterization of iPSC-Derived Neurons.” bioRxiv. https://doi.org/10.1101/2021.11.24.469921.

Rizvi, Abbas H., Pablo G. Camara, Elena K. Kandror, Thomas J. Roberts, Ira Schieren, Tom Maniatis, and Raul Rabadan. 2017. “Single-Cell Topological RNA-Seq Analysis Reveals Insights into Cellular Differentiation and Development.” Nature Biotechnology 35 (6): 551–60.

Satone, V. K., R. Kaur, A. Dadu, H. Leonard, and H. Iwaki. 2020. “Predicting Alzheimer’s Disease Progression Trajectory and Clinical Subtypes Using Machine Learning.” bioRxiv. https://www.biorxiv.org/content/10.1101/792432.abstract.

Sharma, Ankur, Elaine Yiqun Cao, Vibhor Kumar, Xiaoqian Zhang, Hui Sun Leong, Angeline Mei Lin Wong, Neeraja Ramakrishnan, et al. 2018. “Longitudinal Single-Cell RNA Sequencing of Patient-Derived Primary Cells Reveals Drug-Induced Infidelity in Stem Cell Hierarchy.” Nature Communications 9 (1): 4931.

Singh, Gurjeet, Facundo Mémoli, Gunnar E. Carlsson, and Others. 2007. “Topological Methods for the Analysis of High Dimensional Data Sets and 3d Object Recognition.” PBG@ Eurographics 2. http://diglib.eg.org/bitstream/handle/10.2312/SPBG.SPBG07.091-100/091-100.pdf?sequence=1&isAllowed=y.

Strunz, Maximilian, Lukas M. Simon, Meshal Ansari, Jaymin J. Kathiriya, Ilias Angelidis, Christoph H. Mayr, George Tsidiridis, et al. 2020. “Alveolar Regeneration through a Krt8+ Transitional Stem Cell State That Persists in Human Lung Fibrosis.” Nature Communications 11 (1): 3559.

Tustison, Nicholas J., Philip A. Cook, Andrew J. Holbrook, Hans J. Johnson, John Muschelli, Gabriel A. Devenyi, Jeffrey T. Duda, et al. 2021. “The ANTsX Ecosystem for Quantitative Biological and Medical Imaging.” Scientific Reports 11 (1): 9068.

